# Nanojunctions: Specificity of Ca^2+^ signalling requires nano-scale architecture of intracellular membrane contact sites

**DOI:** 10.1101/2023.02.17.528983

**Authors:** Nicola Fameli, Cornelis van Breemen, Klaus Groschner

## Abstract

Specificity of control over virtually all essential cellular functions by Ca^2+^ is based on the existence of separated, autonomic signaling modules. Spatiotemporal definition of Ca^2+^ signals involves the assembly of signaling complexes within the nano-architecture of contact sites between the sarco/endoplasmic (SR/ER) reticulum and the plasma membrane (PM). While the requirement of precise spatial assembly and positioning of the junctional signaling elements is well documented, the role of the nano-scale membrane architecture itself, as an ion reflecting confinement of the signaling unit, remains as yet elusive. Utilizing the NCX1/SERCA2-mediated ER Ca^2+^ refilling process as a junctional signalling paradigm, we provide here the first evidence for an indispensable cellular function of the junctional membrane architecture. Our stochastic modeling approach demonstrates that junctional ER Ca^2+^refilling operates exclusively at nano-scale membrane spacing, with a strong inverse relationship between junctional width and signaling efficiency. Our model predicts a breakdown of junctional Ca^2+^ signaling with loss of reflecting membrane confinement, irrespective of the spatial positioning of the molecular signaling elements. Alterations in the molecular and nano-scale membrane organization at organelle-PM contacts are suggested as new concept in pathophysiology.

## Introduction

Spatio-temporal variations of ionic calcium (Ca^2+^) concentration in the cytoplasm, [Ca^2+^]_i_, referred to as Ca^2+^ signals, regulate the majority of processes that govern cell state and function. This immediately raises the crucial question: “How does an ubiquitous signalling ion, like Ca^2+^, selectively activate each specific function, without triggering all other Ca^2+^sensitive enzymes and ionic transporters?”

Current consensus holds that such selective Ca^2+^ signalling is mediated by specialized junctions between regions of the endoplasmic reticulum (ER, the main intracellular Ca^2+^ store and cytoplasmic Ca^2+^ distribution organelle [1]) and the plasma membrane (PM) or membranes of other organelles. These intracellular compartments form composite signalling elements featuring two closely apposed bio-membranes, endowed with both Ca^2+^ sources and Ca^2+^ receptors, spaced at distances below 50 nm and extending laterally up to hundreds of nm [2]. In 1957 Porter and Palade first described nano-scale junctions [3], between the T-tubule and sarcoplasmic reticulum (SR) in skeletal muscle (triads) and proposed that they played a role in excitation-contraction coupling. Since then similar narrow junctions between cytoplasmic membranes, herein referred to as nano-junctions (NJs), have been identified in every cell type investigated, including endothelium, smooth muscle, myocardium, neurones, T-cells and platelets. Each of these types of NJs employ specific mechanisms to initiate different functions (see table 1). However, the common principle of a large variety of NJs is that the ultrastructure of the nano-space and the composition of the membranes confining this junctional space determine the success of a local Ca^2+^ supply reaching its target.

**Table 1:** Examples of Nanojunctions.

| Junction type | Source | Target | Tissue/Cell Type | Reference |
| --- | --- | --- | --- | --- |
| PM-SR/ER | rNCX* in PM | SERCA2 in SR/ER | smooth muscle,<br>endothelium | [4] |
|  | Orai1 in PM | SERCA2 in ER | endothelium | [5]; [6]; [7] |
|  | TRPC6 in PM | NCX in PM | smooth muscle | [8]; [9] |
|  | RyR2 in ER | NCX in PM | endothelium | [10] |
|  | IP<sub>3</sub>R/RyR in SR | NCX in PM | smooth muscle | [11]; [12] |
| mitochondria-ER | IP<sub>3</sub>R in ER | Ca<sup>2+</sup> uniporter in IMM | endothelium | [13]; [14] |
|  | piezo1 in PM | TRPV1 in PM | acinar cells | [15] |
| lysosome-ER | TPC2 in lysosome | RyR3 in SR | smooth muscle | [16]; [17]; [18] |
|  | TPC2 in lysosome | SERCA (& RyR2?) in SR | heart | [19]; [20] |
| PM-DTS<sup>†</sup> | rNCX in PM | IP<sub>3</sub>R in DTS | platelets | [21] |

\*rNCX = Ca<sup>2+</sup>-entry-mode NCX
<sup>†</sup>DTS stands for Dense Tubular System, equivalent to the smooth ER in platelets

The three principal requirements for site and function specific Ca^2+^ signalling are:

1. co-localization of the source, usually a Ca^2+^ channel, and the targeted Ca^2+^ receptor, such as calmodulin or troponin C or a Ca^2+^ pump
2. the simultaneous presence of Ca^2+^ sources and sinks within a membrane-limited nano-space
3. restricted diffusion between the nano-space and the bulk cytoplasm.

Importantly, these principles apply to any communication between a Ca^2+^ source and its target, even when membrane bridging structures are not involved. Nonetheless, the underlying mechanisms are ill defined because the nano-scale dimensions have thus far defied direct measurements of Ca^2+^ dynamics by confocal, or even super-resolution, microscopy.

Here we set out to analyze the principles underlying the generation of productive Ca^2+^ signals, which are precisely restricted in space and time within membrane confined NJs to enable targeted activation of Ca^2+^ receptors, while preventing spread into the bulk cytoplasm. We set out to test the hypothesis that the presence of “reflecting/confining” membranes within a critical nano-scale distance of the “source membrane” is indispensable for the nano-junction to operate as the critical unit for cell signalling as shown for a wide range of cell types [22]. For our study, we selected the NJ responsible for ER refilling in, among others, vascular endothelial and smooth muscle cells, for which there is consensus that it depends on linked communication between the NCX and SERCA [6]; [7]; [24]; [25]. The ER/SR-PM junction to be modelled here has been reported to refill the SR of vascular smooth muscle in order to sustain asynchronous Ca^2+^ oscillations supporting tonic contraction [26]; [4]. Evidence shows that neurotransmitters stimulate PLC to generate oscillating SR Ca^2+^ release from the SR, while simultaneous activation of TRPC6 supplies Na^+^ to switch the NCX1 into Ca^2+^ entry mode [4]; [9]. Initial simulations using a combination of ultrastructural and physiological data confirmed that linking Ca^2+^ entry via NCX1 to SERCA2 in the superficial SR constitutes a plausible mechanism for refilling smooth muscle SR [24]. An analogous mechanism involving ER-PM junctions in endothelial cells has been shown to participate, in parallel with the action of the STIM-Orai system, in the refilling of the ER after histamine-induced ER Ca^2+^ release [7]; [5].

In order to characterize the operation of typical ER (or SR)-PM junctions with a satisfactorily high level of spatio-temporal resolution, three-dimensional architectural renderings and a stochastic Brownian motion simulator by random walk were used to re-create intracellular organellar membranes, implement known reaction parameters, and analyze the resulting concentration profiles at various membrane separations.

Our results provide a first demonstration that precise positioning of the reflecting membrane of NJs is by itself an essential prerequisite for effective junctional signal transduction.

## Methods

### Preamble and “lay of the land”

The schematics in figure 1 depict the typical spatial organization of ER-PM nanojunctions, as it stems from numerous ultrastructural studies in a variety of cell systems mentioned in the introduction (figure 1A,B). In panel C of figure 1, we report a pseudo-three-dimensional representation of a nanojunctional environment as a set of software objects used in the stochastic simulations comprising the relevant transporters underlying the ER Ca^2+^ refilling mechanism that we address in this article. Referring to figure 1CD, Ca^2+^ ingress into the NJ happens via the Ca^2+^-influx mode NCX1 (blue objects) positioned both in the junctional and extrajunctional PM (purple plane). We have implemented a higher surface density of NCX1 on the junctional PM (450 NCX/μm^2^) than on the extra-junctional PM (18 NCX/μm^2^), and a SERCA2 (green objects) surface density on the junctional ER membrane of 2800 SERCA/μm^2^, in accordance with earlier reports [24]. The typical observed gap width between junctional PM and junctional ER membranes is approximately 30 nm, but has been observed to be considerably narrower for certain junctions. Consistently, junctional membrane spacing is one of the parameter we altered in order to study the effect of ultrastructural disruption on the efficiency of the nanojunction. The lateral extension of the junction was held constant at 400 nm (figure 1C).

**Figure 1:**
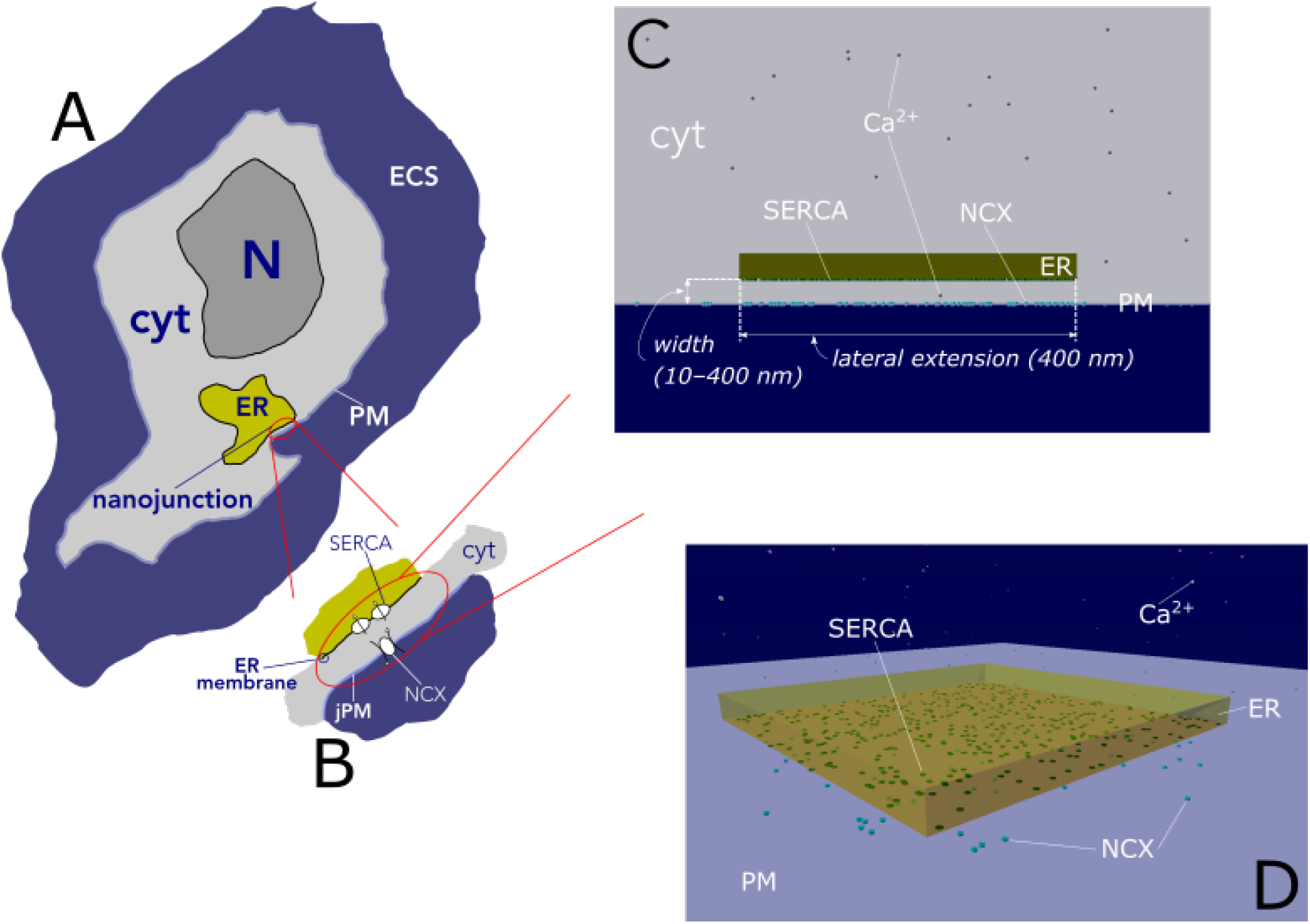
Schematic depiction of the typical location of a cytoplasmic NJ within a cell (purple outline with white background in panel A), in which ER refilling is achieved by Ca^2+^ entry via NCX1 on the junctional PM (jPM, panel B, PM in panels C and D) and Ca^2+^ capture by SERCA pumps localized on the junctional ER membrane (panels B, C and D). The 3D model junction used in the stochastic simulations is shown in panels C (edge-on view) and D (pseudo 3D). The NJ depicted in D is visualized from inside the cell, with the inner PM surface in purple, and the ER in green. The NJ dimensions (width and lateral extension in panel C) and transporter densities are inferred from 2D electron micrographs and from immunocytochemical studies, respectively. N = nucleus; ECS = extracellular space; cyt = cytoplasm.

### Quantitative Model

We developed our quantitative model using the Monte Carlo particle simulation platform MCell/CellBlender (freely available at mcell.org) [27]; [28]; [29], which we have successfully employed for quantitative models in analogous studies before [1]; [18]. In essence, the modelling approach consists of the following main steps:

1. the design of 3D software mesh objects (nets of interconnected triangles by which surfaces can be represented in computer graphics) representing a stylized ER-PM contact region according to typically observed dimensions of the separation between the membranes (around or less than 30 nm) and with the lateral extension of the nano-junction (around 400 nm). These objects are based on electron microscopic ultrastructural characterizations of these junctions as observed in earlier studies [7]; [24];
2. the positioning of the relevant transporters on their respective membranes (NCX1 on the junctional PM (jPM), SERCA2 on the junctional ER (jER) membrane) according to information gathered from the literature on their typical membrane densities and the implementation of the known transporter kinetics ([24] and references therein) and multi-state models, and of the Ca^2+^ diffusivities [32];
3. the simulation of molecular Brownian motion in the cytosol by random walk algorithms; this phase is performed by “number crunching” component of the stochastic particle simulator MCell. This reproduces the randomness of the molecular trajectories, of the ion transporter flickering and of the relevant chemical reactions by probabilistic algorithms governed by pseudo-random number generators. This enables the simulation of a number of microphysiological processes, all stochastically different from one another. The average outcome of the processes, to mimic the instrumental output during experimental measurements, is obtained by taking the mean (or median) of a desired quantity, e.g., Ca^2+^ captured by SERCA2, over a large number of simulations all initiated with different random number generator seeds.

## Results

### Junctional [Ca^2+^] quickly reaches steady-state with only few, freely diffusing Ca^2+^

In a first set of simulations, we aimed to understand the timescale with which our basic ER-PM junction system would reach steady-state in terms of junctional Ca^2+^ concentration, [Ca^2+^]_NJ_. Having set up the system with the geometry described in figure 1, with a membrane separation of 30 nm and the transporter densities and reaction kinematics properties reported above, we ran 100 simulations in each of which Ca^2+^ was released into the junction by NCX1 and was made to random-walk until it either was captured by a SERCA2 pump or exited the junction. At each time step, simulations measure and record the number of Ca^2+^ present in the NJ in an output file. In each separate simulation, the positions of the NCX1 and SERCA2 sets were randomized so as to minimize artifacts from particular positioning of transporters on the junctional membranes. Since the known NCX1 turnover rate is about 3500 Ca^2+^/s, we expect a low number of Ca^2+^ to populate the NJ at any given time at sub-millisecond timescales. In figure 2A, we report the mean and standard error of the “measured” [Ca^2+^]_NJ_ temporal profile (the [Ca^2+^]_NJ_ in moles/L is calculated by dividing the recorded number of Ca^2+^ in the NJ by the volume of the NJ and by Avogadro’s number). Figure 2B, illustrates the actual number of Ca^2+^ in the NJ *vs* time over a simulation time of 0.5 ms in a separate set of 100 simulations.

**Figure 2:**
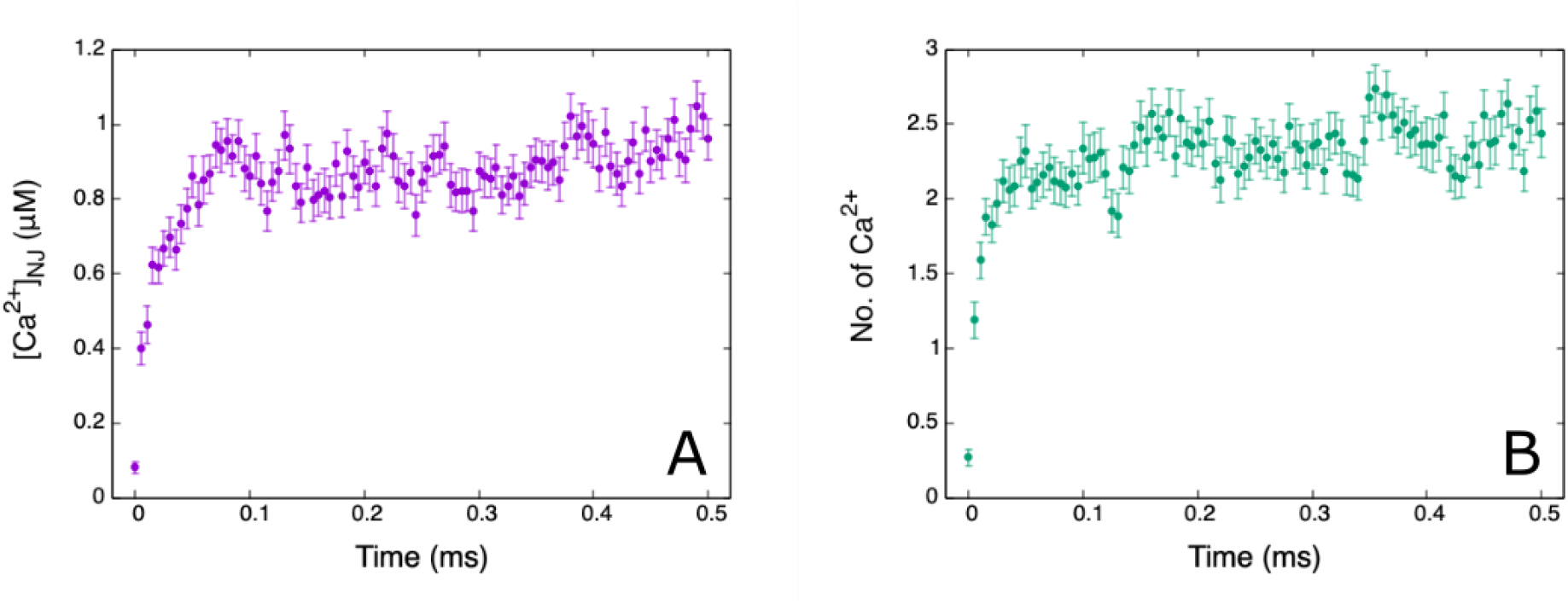
A: [Ca^2+^]_NJ_ temporal profile in a typical model junction during a 0.5 ms interval. The junctional membrane separation is 30 nm and the lateral extension is 400 nm × 400 nm (see figure 1D). B: Number of Ca^2+^ in the same model junction during the same interval. Reported values are mean ± SE (n = 100).

### Junctional width (membrane separation distance) is a crucial determinant of NCX1-to-SERCA2 Ca^2+^ transfer

Given the paucity of Ca^2+^ in this junctional system, the effectiveness of the nanojunctional communication must rely on highly efficient transfer of Ca^2+^ from source (NCX1) to sink (SERCA2). In a set of simulations, we counted the number of times the reaction SERCA2 + Ca^2+^*→* Ca.SERCA2 (that is SERCA2 with one bound Ca^2+^) took place in the same time frame within which the system reaches [Ca^2+^]_NJ_ steadystate (determined in figure 2) as a function of the membrane-membrane separation distance (each simulated configuration differs from the others simply by translating the ER software object a certain distance away from the PM object in a direction perpendicular to the latter). In figure 3A, we report the results from 6 separation distances, from 10 nm to 400 nm, with each of the points being the mean ± SE over 100 simulations. The very wide 400-nm separation case was implemented to investigate the efficiency of essentially bulk Ca^2+^ transport between NCX1 and SERCA2. Figure 3B illustrates the time course of fractional Ca^2+^ transfer from NCX1 to SERCA2. Both sets of results show a dramatic drop in junctional performance with increasing gap width and the virtual loss of function at a membrane spacing of 400 nm.

**Figure 3:**
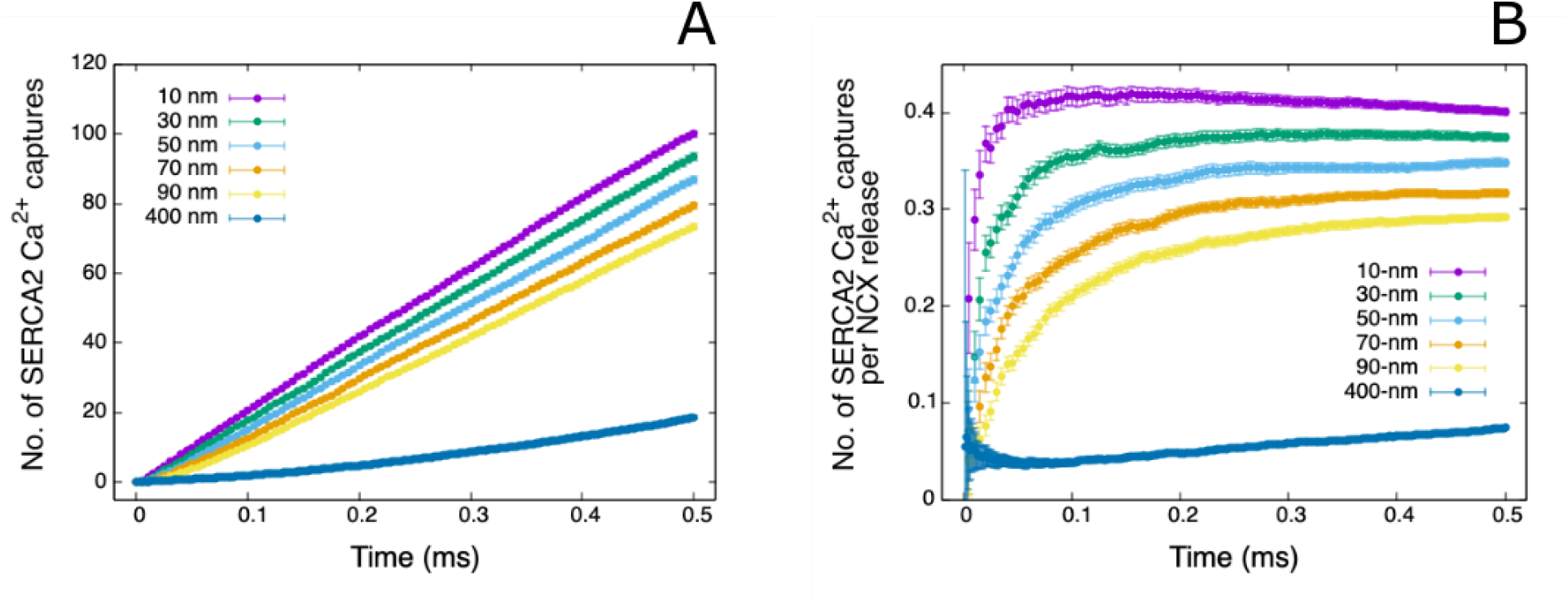
A: Cumulative number of SERCA2 Ca^2+^ capture events vs time during a 0.5-ms NCX1 Ca^2+^ entry period. B: Fractional Ca^2+^ capture events by SERCA2 vs time. Simulations were run for 6 different junctional widths, as indicated in the legend. Each point is the mean ± SE from 100 simulations; note that the error bars can be roughly the size of the symbols.

### Reflective membrane confinement of the junctional space is essential for Ca^2+^ transfer

To further elucidate the role of the very presence of the membranes as confining elements to assist and facilitate Ca^2+^ signalling and junctional performance, and ultimately cell function, we also ran sets of simulations, in which we made the jER membrane transparent to Ca^2+^, but left the SERCA2 pumps in place, distributed as in the “regular” NJ, that is on a plane a given distance away from the jPM as illustrated in the right-side panels in figure 4. In practice, a comparable scenario might be generated when the junctional contact area is strongly reduced in lateral extension with minimal contribution of the ER membrane. Indeed, dynamic changes in PM-ER junctional contact area have been reported [33]; [34]; [35]. We therefore characterized the consequences of simply lacking the ER membrane confinement. In this configuration Ca^2+^ arriving at the jER will not be reflected back into the junction, but will continue their random walk through.

**Figure 4:**
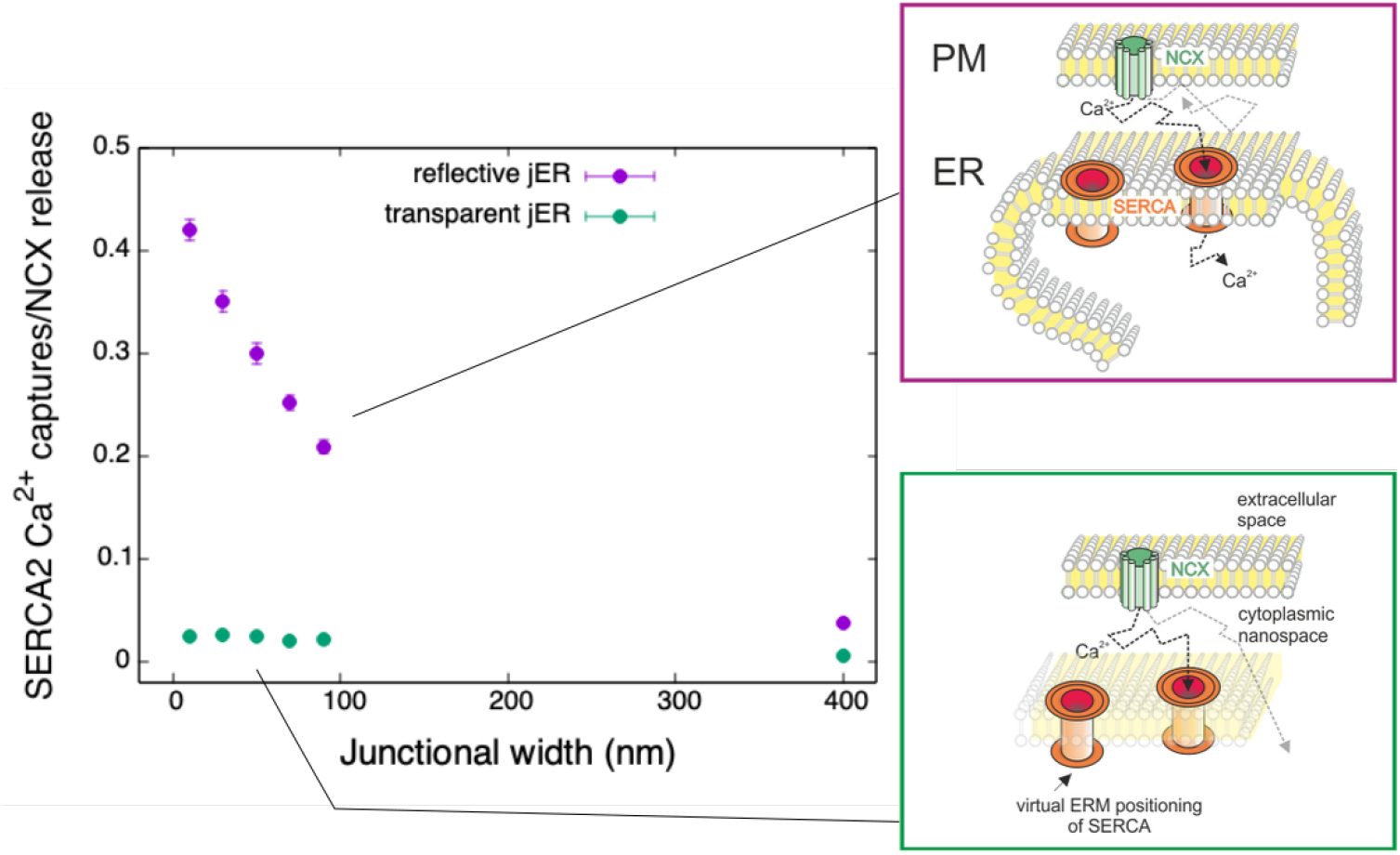
Left: Number of SERCA2 Ca^2+^ capture events after a 0.1 ms NCX1 Ca^2+^ release period from the jPM as a function of junctional width in two scenarios: one with a “Ca^2+^-reflective” (purple data and top right panel) and another without (or “Ca^2+^-transparent”; green data and bottom right panel). The Ca^2+^ source and sink molecules remain in the same position under the two computational tests with the only difference being the presence or absence of confinement of the junctional space by the ER membrane. Broken lines in the diagrams on the right represent 3D random walk trajectories. ERM = ER membrane.

Again, we changed the distance of the SERCA2 plane from the junctional PM to study the effect of this separation on SERCA2 capture efficiency. We report the results in figure 4, left-side panel. These results demonstrate a dramatic impact of the lipid bilayer architecture of Ca^2+^ signaling NJs independent of spatial positioning of Ca^2+^ source and sink or target. Suitable apposition of the junctional membranes along with sufficient lateral extension of the ER membrane confinement are shown to be prerequisites for efficient NJ communication.

### Confinement of the junctional space enables the high frequency of Ca^2+^ hits at the ER required for efficient SERCA2-mediated Ca^2+^ uptake

Since we can expect that on average 3 NCX1-released free Ca^2+^ will populate a NJ at any given time (figure 2B), in separate sets of simulations we let 3 Ca^2+^ random walk their way inside a junction from positions near the junctional PM and counted how many times each ion would hit the jER or the jPM before being captured by a SERCA2 pump. The calculated median hit count for each ion over 100 simulations was then reported as the mean of the medians over 3 ions in figure 5AB. We also deemed it relevant to report the dramatically increased time scales at which Ca^2+^ captures take place when junctional width is extended (figure 5C).

**Figure 5:**
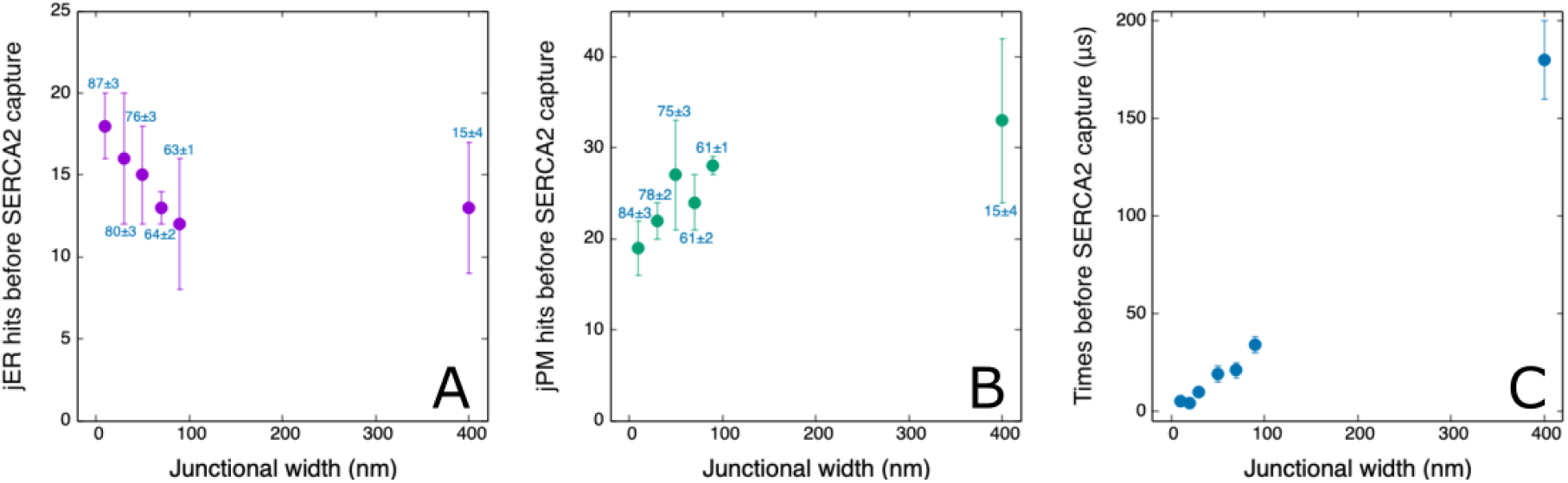
A: Mean ± SD of number of times a Ca^2+^ hits the junctional ER surface before being captured by a SERCA2 pump as a function of junctional width. B: As in A, but counting junctional PM hits. The numbers in blue indicate the mean percentage probability (±SD) that a released Ca^2+^ will get captured at each width (the remainder represents the probability that ions escape the junction). C: Mean ± SD times before SERCA2 capture vs junctional width.

### ER membrane-delimited 2D-RW has the potential to remarkably accelerate junctional Ca^2+^ transfer

We have previously considered the possibility that part of the Ca^2+^ trajectories in NJs involve ion exchange diffusion on the surface of the peripheral ER after the first hit [36]. Figure 6 illustrates results obtained by extending our model system (a model junction with a 30-nm width and 400-nm lateral extension) to include a process of 2D-surface diffusion of the Ca^2+^ ions, either as an exclusive mechanism to reach its SERCA2 target subsequently to a first hit at the ER membrane or as an additional trajectory option alongside successful 3D-RW trajectories. Our simulations predict that irrespectively of allowing successful 3D-RW Ca^2+^ transfer, the 2D-RW trajectory option enhances efficiency of junctional operation in terms of Ca^2+^ capture/transfer rates. These findings strongly suggest the need to refine the current picture of junctional Ca^2+^ signalling, not only by including the aspects of membrane nano-scale architecture and reflector-type confinement within the junctional signalling space, but also taking into consideration the concepts of Ca^2+^ binding to phospholipids and 2D diffusion of Ca^2+^ on the surfaces of the confining membranes as discussed below.

**Figure 6:**
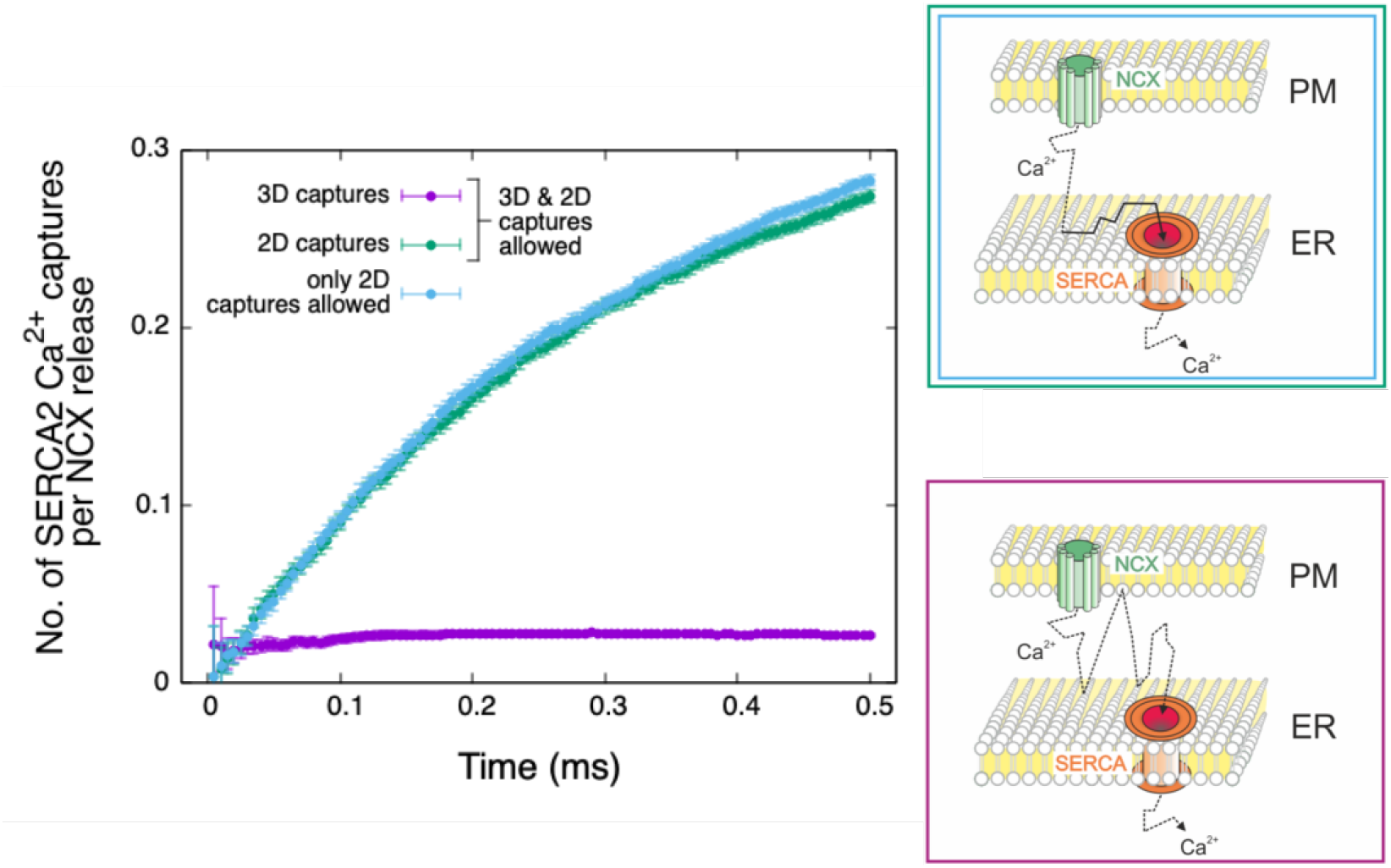
Comparison of the effects of Ca^2+^ only allowed to do 3D-RW vs 3D-RW until the first hit on jER, followed by 2D surface diffusion with the same diffusivity as the 3D random-walking ions. Purple and green data are derived from simulations allowing for captures from both types of RW; light blue data are derived from simulations allowing captures from 2D-RW only. The model junction employed has a membrane separation of 30 nm and lateral extension of 400 nm × 400 nm.

## Discussion

### Novel insights into the nature of Ca^2+^ trajectories in NJs

The main question we are addressing is: What is the role of an ion-confining membrane in the activation of junctional (low-affinity) Ca^2+^ receptors? The results of our stochastic simulations of individual Ca^2+^ trajectories within the NJ, which links NCX1 and SERCA2 pumps, point to a fundamental role of such junctional membranes and raises important questions related to the architecture of the junctional nanospace harbouring both the Ca^2+^ source and its target.

### The need for stochastic simulations

In many instances in the literature on specific Ca^2+^ receptor targeting one relies on the idea of “microdomains” as the localized high-[Ca^2+^] volume near the mouth of a transporter, within which a target must be placed for efficient activation. However, if we accept, as is supported by numerous reports (see table 1), that membrane junctions are Ca^2+^ signalling structures of sub-micron dimensions, we must also revisit the idea of ion concentration within the junctional spaces. This is especially important when lower capacity transporters are involved in the signalling mechanisms, like for example in the NCX1-SERCA2 Ca^2+^ communication in SR/ER refilling in smooth muscle and endothelial cells, or in the Orai-SERCA2 phase of SOCE in many cell types. Both NCX and Orai have Ca^2+^ capacities of the order of 4000 ions/s, which yield a junctional “population” of about three Ca^2+^/junction at any given time, using the typical free Ca^2+^ diffusivity measured in cytosol [32].

The results reported in figure 2A show that during activation of the Ca^2+^-entry-mode of NCX, the [Ca^2+^]_NJ_ rises from 0.08 M to 1.0 M, in a representative model NJ containing NCX1 and SERCA2 densities expected in smooth muscle and endothelial cells [24]; [37] and dimensionally within the typically observed range. Taken at face value, this rise in concentration is sufficiently above the Ca^2+^ affinity of the target SERCA2. However, figure 2B also shows that this rise in [Ca^2+^]_NJ_ results from an increase in the number of Ca^2+^ in the junctional nano-space from 0.2 to 2.5. This low number of Ca^2+^ per junction renders physically meaningless the above-mentioned concept of Ca^2+^ signaling within microdomains and is in many aspects incompatible with the application of modeling approaches based on reaction-diffusion equations, which rely on averages over large numbers of ions. Based on these observations, we employed a stochastic simulation approach to recreate the Ca^2+^ RW trajectories within dimensionally and biophysically accurate model NJs, to provide information that cannot be gained by conventional diffusion equations, nor, at this time and at this scale, by fluorescent Ca^2+^ indicators.

### Efficient NXC-SERCA2-mediated Ca^2+^ transfer requires a junctional width of less than 50 nm

Our modelling approach provides unambiguous support for the concept of NJ Ca^2+^ signalling requiring close apposition of the junctional membranes with gap widths being a critical determinant of signalling efficiency. Simulations of Ca^2+^ transport from NCX1 as a PM-resident source to the ER lumen via SERCA2 as an ER-resident target unveiled an inverse relation between gap width and Ca^2+^ transport rate. The number of Ca^2+^ ions captured by SERCA2 within a given period of time dropped dramatically with increasing the width of the junctional gap (Figure 3 and 4). Transport rates became inefficient when the gap width exceeded 50 nm. This result is in line with previous studies analyzing the relation between membrane ultrastructure and Ca^2+^ signalling at ER-PM junctions. Disruption of the junctional membrane architecture and specifically interventions that separate the PM from ER abolish NJ signalling. This was convincingly shown for vascular smooth muscle by Lee et al. [38], who provided EM data to document calyculin-A-induced NJ membrane separation. Calyculin-A treatment did not affect the ability of the SR to accumulate and release Ca^2+^, but did abolish the asynchronized Ca^2+^ oscillations and tonic contraction in response to alpha-adrenergic stimulation, both of which depend on direct, junctional refilling of the ER/SR. Pritchard et al [39] also showed that microtubules play an essential role in maintaining the narrow distancing between PM and SR at the NJs involved in RyR-BKCa coupling. Nocodazole caused a 4-fold widening of the NJs, from approximately 20 nm to 80 nm, which uncoupled the sparks from hyperpolarization.

Precise spatial alignment of ER-PM junctional membranes by a wide range of tethering complexes has long been recognized as indispensable for the cellular functions of these contact sites including their role in lipid homeostasis, control of membrane structures but also Ca^2+^ signalling [40]; [41]; [42]. Importantly, a junctional gap even below 20 nm is required for ER Ca^2+^ refilling by the STIM/Orai Ca^2+^ entry machinery, as this highly specialized signalling process requires molecular bridging of the ER-PM junctional gap [33]; [43]. For this particular Ca^2+^ signalling scenario, strategic positioning of the signalling molecules is essential, and a complete loss of function is inevitable when the junctional membranes are separated beyond a critical distance. By clear contrast, the junctional signalling process modeled in our study is different in nature. NCX1/SERCA2-mediated junctional Ca^2+^ transfer does not involve dynamic physical coupling of the junctional membranes and therefore lacks the requirement for strategic positioning of source and target proteins. This is clearly evident from our simulation of the NCX1/SERCA2-mediated junctional Ca^2+^ transfer, featuring an ER membrane structure without barrier function for Ca^2+^(“transparent” membrane). Despite retaining an adequately close positioning of the signalling molecules, mere elimination of the ER membrane barrier and consequently of the impact of this spatial confinement on 3D-RW of junctional Ca^2+^, dramatically reduces signalling efficiency (Figure 3 and 4) to a level typically observed when junctional membranes are separated by 400 nm or more. Hence, we provide the first demonstration for the importance of junctional gap width and of the NJ membrane confinement for signalling performance, besides the widely recognized aspects of spatial organization of signalling proteins. As a mechanistic basis of this observation, our NJ model suggests that successful junctional transfer of a Ca^2+^ ion by 3D-RW within the NJ gap involves a fairly constant number of “nonproductive” hits at NJ confining membranes before its capture by SERCA2 (Figure 5AB). Therefore, the time required for productive trajectories increases steeply with the gap width (Figure 5C), resulting in the drop in NJ Ca^2+^capture rates and loss of signaling function presumably at gap widths larger 50 nm.

### Ca^2+^ binding to and coordination at junctional phospholipid (PL) membrane surfaces is likely to promote signalling efficency

The above analysis is based on 3D-RW simulations within a simplified architecture incorporating inert reflecting surfaces. However, electron microscopy of NJ ultrastructure and biochemistry of PL membranes indicate that this picture is far from complete. In reality the surfaces limiting the nano-space are highly irregular and composed of various charged PL head groups and neutral cholesterol molecules with interspersed membrane proteins. At rest the junction is inhabited by fewer than one hydrated freely diffusible Ca^2+^, which rises at the peak of activation to approximately one hundred Ca^2+^ ions (i. e., 30 μM). The polar head-groups of PLs facing the cytoplasm contain several negatively charged oxygen groups each, which means that with an average single PL surface area of 0.64 nm^2^ [44] the reflecting membrane surfaces lining each NJ contain more than 100,000 negative binding sites. In other words, the relevant stoichiometry within an NJ is that there are three to four orders of magnitude greater numbers of negative binding sites than there are Ca^2+^ ions. Although direct measurements are lacking, these numerical considerations lend credence to reports, which estimate that as much as 90 to 99% of Ca^2+^ residing in narrow inter-membrane, cytoplasmic spaces is bound. Hence, it is plausible to assume that a prominent fraction of the Ca^2+^ involved in NJ signalling is bound to PL head groups [45]. Computational modeling of interactions between divalent cations and phospholipid-containing biomembranes demonstrated that Ca^2+^ ions indeed desolvate readily to bind to and bridge multiple PL headgroups at membrane surfaces, thereby generating a two-dimensional network with Ca^2+^ residing within PL clusters. This process, which has been extensively studied for phosphoinositides (PIP_2_) occurs even at low micromolar divalent concentrations and Ca^2+^ clearly outperforms Mg^2+^ in terms of its PL bridging function [46]; [47]. Hence, it appears reasonable to picture a scenario in which Ca^2+^ entry into the NJ drives reversible Ca^2+^ loading onto the NJ confining membranes associated with displacement of other counterions such as Mg^2+^ at negatively charged PL headgroups. Although direct evidence is lacking, it is plausible that the NJ membranes function as a buffer system for Ca^2+^ and are subject to phasic accumulation of Ca^2+^ within the PL network surrounding integral membrane proteins such as NCX1 and SERCA2. The lipid nano-environment of signalling proteins, recently designated as the “functional lipidome” [48], therefore needs consideration as a pool of Ca^2+^ ions that is potentially relevant for signalling by Ca^2+^ receptors or transporters. Specifically for SERCA, the Ca^2+^ pool represented by its functional phospholipidome is the most likely source from which Ca^2+^ ions enter the transporters via peripheral, hydrophilic entry pathways, formed by rows of carbonyl oxygens which guide the divalent Ca^2+^ by a process of serial ionic binding towards the key Ca^2+^ translocation sites in the transmembrane domain [49]. This mechanism is in line with our simulation of the 3D-RW NJ model, which has a minimum size of 20 nm^2^ for the functional Ca^2+^ target site, which is substantially larger than a single divalent coordination site. The concept of surface-delimited NJ transport of Ca^2+^ may be extended by considering that a major part of the junctional Ca^2+^ transfer involves 2D-RW along the NJ-confining membranes based on the interfacial ion exchange diffusion concept proposed recently [36]. Several lines of evidence support the above hypothesis and warrant further investigation. Lateral diffusion of both Ca^2+^ and Na^+^ on PL surfaces has been demonstrated in artificial membranes [51]; [52]; [53]. The mechanism proposed was one of serial ionic bonding to negative PL headgroups lining long narrow pores in a millipore filter followed by removal of bound Ca^2+^ on the trans side of the membrane distal site by chelation [52]. Lateral exchange diffusion on the junctional SR/ER surface may involve a similar mechanism of serial reversible binding to negatively charged PL head-groups, followed by Ca^2+^ sequestration by SERCA generating a renewable supply of target binding sites [52]. Although the rate of 2D ion exchange diffusion may be slower than 3D diffusion in the cytoplasm, the fifteen fold increase in the SERCA2 capture rate shown in figure 6 supports a role for 2D surface diffusion in NJ Ca^2+^ signalling. An additional advantage of 2D Ca^2+^ exchange diffusion on the ER surface would be access to SERCA2 outside the junctional domain and thereby increasing the efficiency of SR/ER refilling.

The results of our stochastic modelling demonstrates the superior efficiency of a Ca^2+^ signalling process that includes 2D random walk components as compared to a simple 3D-RW model (figure 6). It appears therefore attractive to speculate that NJ Ca^2+^ signalling involves for a large part 2D interfacial exchange diffusion. This hypothesis is in line with observations indicating a striking role of the membrane lipid composition as a determinant of NJ signalling efficiency. In this context it is of note that particularly fast-operating, highfrequency muscle organs have been found to strictly require a certain membrane phospholipid composition which is greatly enriched in docosahexanoic acid [54].

### Physiological and Pathological relevance

Since the trajectory of Ca^2+^ from source to target within the NJ is central to site and function specific Ca^2+^ signaling in all animals, it is important to understand how it may affect growth and development, aging, health and disease. However, it would take a major review to cover the health implications of all site and function specific Ca^2+^ signaling and therefore only examples related to our model of the NJ, mediating privileged transfer of Ca^2+^ between the extracellular space and ER/SR lumen, will be alluded to.

In order for the ER to regulate the Ca^2+^ fluctuations in various nano-spaces between its outer membrane and the membranes of other organelles, including the PM, independently from the bulk cytoplasm, it requires direct routes for unloading and refilling to and from the extra cellular space [1]. Chronic diseases can disrupt this homeostatic mechanism and lead to ER Stress, involving loss of ER Ca^2+^ content and accumulation of unfolded proteins in the ER lumen. The effect of diminished ER Ca^2+^ content has been observed in vascular smooth muscle of a line of mice, harbouring the fibrillin defect characteristic of Marfan Disease Syndrome in humans, in terms of a lower frequency of asynchronous calcium oscillations and reduced contractile force [55]. The Ca^2+^-entry mode NCX, within the type of NJ referred to above, could even be a promising therapeutic target site for a number of chronic diseases. Piccialli et al. [56] showed that upregulation of Ca^2+^-entry-mode NCX in a mouse model of Alzheimer’s disease promotes ER refilling in primary neurones to protect against ER Stress and death.

Finally a role of NJ decay in aging has been demonstrated in vascular smooth muscle of healthy mice, by parallel losses of close contacts between the SR and PM, as visualized by electron microscopy, and asynchronous Ca^2+^ oscillations recorded by fluorescence microscopy [57]; [58].

## Conclusion

By stochastic modeling of Ca^2+^ trajectories within ER-PM nanojunctions, we identify as yet unrecognized structural requirements for efficiency and specificity of cellular Ca^2+^ signaling. Our results suggest that both the nano-scale dimensions of the junctional membrane gap as well as the continuous confinement of junctional contact regions are indispensable for signal transduction. Alterations in these critical structural features of NJs need consideration as a potential basis of human disease and as targets for new therapeutic interventions.

